# Phytoplankton adaptive resilience to climate change collapses in case of extreme events – A modeling study

**DOI:** 10.1101/2023.02.06.527319

**Authors:** Boris Sauterey, Guillaume Le Gland, Pedro Cermeño, Olivier Aumont, Marina Lévy, Sergio M. Vallina

## Abstract

As climate change unravels, ecosystems are facing a warming of the climate and an increase in extreme heat events that are unprecedented in recent geological history. We know very little of the ability of oceanic phytoplankton communities, key players in the regulation of Earth’s climate by the oceans, to adapt to these changes. Quantifying the resilience of phytoplankton communities to environmental stressors by means of adaptive evolution is however crucial to accurately predict the response of marine ecosystems to climate change. In this work, we use an eco-evolutionary model to simulate the adaptive response of marine phytoplankton to temperature changes in an initially temperate oligotrophic water-column. By exploring a wide range of scenarios of phytoplankton adaptive capacity, we find that phytoplankton can adapt to temperature increases –even very large ones– as long as they occur over the time scale of a century. However, when rapid and extreme events of temperature change are considered, the phytoplankton adaptive capacity breaks down in a number of our scenarios in which primary productivity plummets as a result. This suggests that current Earth System Models implicitly assuming perfect and instantaneous phytoplankton adaptation to temperature might be overestimating the phytoplankton’s resilience to climate change.

## I. Introduction

### I.1. Phytoplankton adaptation and climate change

The photosynthetic release of dioxygen and fixation of carbon and other essential elements (N, P, Fe, Si) by marine microbial communities as they produce biomass is a core component of the biogeochemical machinery regulating the chemistry of the atmosphere and oceans, thereby affecting the climate of our planet (Field et al. 1998, Henson et al. 2011). In turn, the oceanic environment (nutrients availability, temperature, pH, etc.) determines the functioning of phytoplankton communities as phytoplankton acclimates and evolutionarily adapts to the spatial and temporal variability of environmental conditions (Litchman et al. 2012). As climate changes, and will keep changing in the next hundreds of years, phytoplankton will face one of the most rapid and diverse environmental shift in the history of our planet (Bopp et al. 2013), combining global secular trends of ocean warming, acidification, stratification and desertification to ever more frequent extreme events (e.g., Gruber et al. 2021, Burger et al. 2022). This raises the questions of the ability of marine phytoplankton to evolutionarily adapt to such changes and of the potential implications regarding the continued role of oceans in climate regulation.

The extent of the phytoplankton ability to adapt to environmental changes is currently unknown, but expected to be substantial. Experimental and *in situ* observations (Padfield et al. 2016, Irwin et al. 2015) tend to suggest that this process can operate in a matter of years. A single liter of oceanic water in the euphotic zone contains 10^6^ to 10^9^ phytoplankton cells (Flombaum et al. 2013) reproducing approximately once a day (Ward et al. 2017). As the environment changes, the large pool of phytoplankton individuals and their rapid generation time allow advantageous mutations to rapidly appear and take over communities through selective sweeps. Yet, will adaptive processes be sufficiently rapid to mitigate the potential deleterious effect of climate change on phytoplankton communities? Given how arduous monitoring evolutionary processes in natural systems is and how little we know about the adaptive properties of the phytoplankton as a result, it is for now difficult to answer this question with certainty and to infer how important phytoplankton adaptation to climate change is for the evolution of ocean biogeochemistry. Yet, what we cannot measure *in situ*, we can model *in silico*. As a first step toward answering these questions, we use an ocean model explicitly accounting for the adaptive evolution of the physiological properties of marine phytoplankton under climate change. We explore a wide range of assumptions regarding the adaptive properties of phytoplankton communities and focus specifically on the adaptive response of the phytoplankton to temperature changes.

### I.2. Phytoplankton and temperature

The response of phytoplankton communities to temperature changes involves two main mechanisms (Eppley 1972, Norberg 2004, Grimaud 2017; Fig. 1A and S1). At the scale of individual cells, growth is typically maximized for a given temperature (the optimal growth temperature, *T_opt_*), above and below which growth drops to zero, the drop being much swifter towards warmer temperatures (see supplementary discussion). The resulting thermal reaction norm represents the thermal niche of an individual phytoplankton cell or, in other words, the physiological ability of this individual to tolerate temperature changes.

**Fig. 1:**
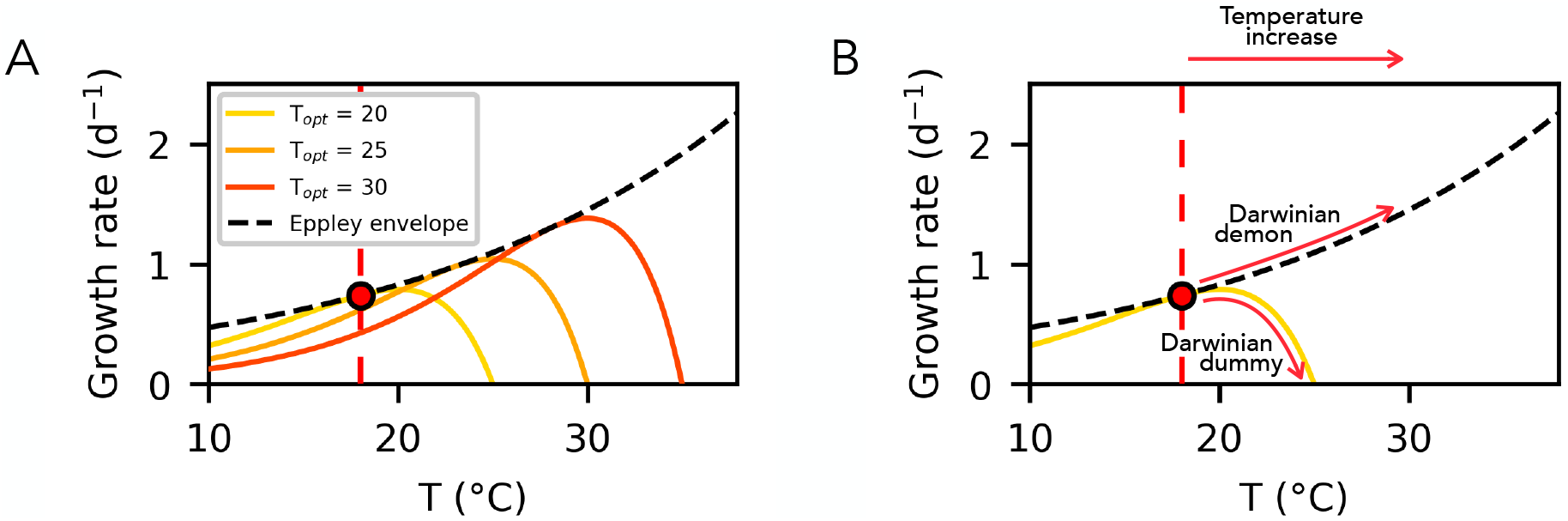
Phytoplankton response to temperature changes. Panel A shows the observed relationship between the growth of phytoplankton individuals and temperature given the thermal niche of those individuals. Each colored thermal reaction norms corresponds to the thermal niche of a specific type of individuals, each niche being characterized by a temperature of optimal growth (*T_opt_*). Panel B shows the expected physiological and evolutionary response of phytoplankton communities to a temperature change.

Every one of these thermal reaction norms can be placed under an increasing power law envelope. This so-called Eppley envelope (from the pioneering work of Richard Eppley, 1972) thus describes the maximum growth rate achievable by any phytoplankton population given a specific temperature. Note that here we use the term population to designate a group of phytoplankton individuals having the same value of the trait *T_opt_*. In a phytoplankton community composed of several competing phytoplankton populations and in the absence of any other selection pressure, one would expect that the phytoplankton population achieving maximal growth given the temperature of the local environment (i.e., the population whose thermal reaction norm is equal, for the temperature of the system, to the Eppley envelope; Fig. 1A) would outcompete the others in the long run. The Eppley envelope therefore corresponds to the optimal evolutionary response of the phytoplankton community to a temperature change.

Depending on the assumption made on whether phytoplankton communities can evolutionarily adapt or not to temperature changes, the predicted change in phytoplankton growth in response to a temperature increase switches from one extreme to the other (Fig. 1B). When assuming that phytoplankton communities can adapt perfectly and instantaneously to temperature fluctuations, the growth rate is expected to increase exponentially with temperature. Henceforward, we will call this scenario the Darwinian demon scenario (from Law 1979). By contrast, when assuming that phytoplankton communities cannot adapt, the phytoplankton response to the temperature change relies solely on its physiology and the growth rate swiftly drops to 0 as temperature increases toward and beyond the tolerance threshold (see supplementary discussion and supplementary table 1). We will call this second scenario the Darwinian dummy scenario.

Which of these two extreme and opposite scenarios is more realistic is expected to depend mostly on the time scale of the temperature change considered. The physiological response of the phytoplankton spans over the lifetime of individuals (few hours to few days, Ward et al. 2017). Evolutionary adaptation on the other hand is a multigenerational process that requires phytoplankton populations carrying newly adapted traits to appear though mutation and to progressively take over the community, replacing through competitive exclusion older, less adapted populations.

Most global-scale models of marine ecosystems assume that the temperature dependence of phytoplankton growth rate follows the Eppley envelope, regardless of the time scale of the temperature change considered (see Henson et al. 2021 as a recent example). In other words, most models consider that the thermal adaptation by phytoplankton is perfect and instantaneous. Interestingly, many of the people using those models are not conscious of this strong implicit assumption. For instance, Henson et al. (2021) state that in their model phytoplankton “do not evolve or adapt to changing conditions” while making that very same implicit assumption.

Is the assumption of perfect thermal adaptation by the phytoplankton fair? The relatively good match between the experimentally evaluated thermal norms of phytoplankton populations and *in situ* temperatures (Thomas et al. 2012, Grimaud et al. 2015) constitutes indirect evidence of the ability of phytoplankton communities to adapt to some extent to temperature. This does not provide, however, information on how rapidly adaptation occurs. Monitoring of shifts in the thermal niche of phytoplankton communities *in situ* and in laboratory experiments of artificial selection suggests that adaptation occurs over a few hundreds of generations (from a few months to few years depending on the taxa considered; Irwin et al. 2015, Padfield et al. 2016). Still, it remains unclear what the actual mechanisms involved in the observed shift in functional composition are; e.g., does it rely on species sorting on a preexisting standing variation or on selection upon *de novo* diversity generated through mutations (Litchman et al. 2012, Collins et al. 2014)? Furthermore, it is unclear whether general evolutionary properties of phytoplankton communities can be inferred from those observations.

Although few ocean ecosystem models explicitly account for the thermal adaptation of the phytoplankton (bur see Grimaud et al. 2015, Demory et al. 2019, Le Gland et al. 2021, Ward et al. 2021; see Ward et al. 2019 for a review of the approaches used to model evolution in ocean ecosystems), none of them addresses the question of the evolutionary response of phytoplankton communities to changing temperatures and of the characteristic time scale of this process. Here we use an eco-evolutionary model explicitly reproducing the functional shift in phytoplankton communities as a result of a mutation-selection process to evaluate the phytoplankton response to long term (secular trend of global warming) and short term (extreme events) temperature changes. We do so in the context of an initially temperate tropical water-column, exploring a wide range of assumptions regarding the adaptive capacity of the phytoplankton in order to address the lack of experimental constraints. The goal is to ask depending on the assumptions of the evolutionary model: How will phytoplankton activity change in response to temperature changes? How reliant on evolutionary adaptation (vs acclimation) is that change? How wrong are we when we assume perfect and instantaneous adaptation?

## II. Methods

### II.1. Model

We use the same setup of 1D model of water-column as in Le Gland et al. 2021, which resolves the vertical physics (vertical mixing by turbulent diffusion) of a temperate subtropical system similar to that of the Bermudas (DuRand et al. 2001, Saba et al. 2010). This marine biogeochemical model resolves the dynamics of several plankton populations (phytoplankton and zooplankton) and of the concentration of dissolved and particulate forms of elements (carbon, nitrogen, phosphorous, silicate, iron) essential to phytoplankton growth. Phytoplankton consumes these elements following a Monod formulation (Monod 1949), and is itself consumed by zooplankton. A recycling term then closes the ecosystem, and the biomass lost to grazing or plankton death is recirculated in the system as dissolved and particulate organic elements (see Le Gland et al. 2021 for details).

The relevance of our model is that, contrary to most ocean ecosystem models, it simulates the variation of the functional composition of the phytoplankton community through time as the result of natural selection. Our approach is based on the SPEAD model (Le Gland et al 2021) that simulates adaptive evolution by means of “trait-diffusion”. Instead of simulating the dynamics of a single phytoplankton population with fixed traits as most models do, we simulate the dynamics of the abundance of 50 ecotypes, *P_i_*, each characterized by a specific thermal niche, i.e., by a specific temperature of optimal growth, *T_opt_*(*i*), ranging from 18 to 50.34°C (Fig. 1; see the supplementary materials for details on the way this thermal niche is parameterized). Each of these ecotypes grows and dies depending on the match of its thermal niche to the environmental temperature. At each division event, a mutation can occur so that the produced offspring ends up with a thermal niche different from that of its ancestor. This process is driven by a key parameter of the model, v, which is the probability of mutation per division or mutation rate (see the supplementary material). This parameter determines the ability of the phytoplankton to generate functional diversity through mutation, a necessary condition to Darwinian evolution. Finally, individuals are moved around within the water column through vertical mixing.

To summarize, the functional composition of the phytoplankton community (i.e., the biomass distribution amongst the 50 functional types) varies as the combined result of three explicitly simulated mechanisms: (*i*) mutations, continuously introducing functional diversity in the community; (*ii*) vertical mixing redistributing functional diversity throughout the column; and (*iii*) competitive sorting selecting within the available functional diversity the ecotypes that are the most adapted to the local environmental conditions. The first two mechanisms tend to increase phenotypic diversity in the community. The third mechanism tends to drive it down. Note that our modeling approach typically results in trait distributions being characterized by very long tails, i.e., many non-viable ecotypes are unrealistically maintained in the system at infinitesimal levels by the mutational process. To prevent this from happening, we add an Allee effect (Allee et al. 1949): below a certain biomass threshold, the biomass specific growth rate of phytoplankton ecotypes steeply decreases with biomass (see the supplementary materials). Such an inverse density dependence of the growth rate is usually linked to the deleterious effects of genetic inbreeding, of demographic stochasticity, and of losses of group advantages as populations get scarce (Courchamp et al. 1999, Fadai et al. 2020).

### II.2. Steady state and regimes of climate change

We study the predictions of the model in two types of environmental context. First, we consider the characteristics of the seasonal equilibrium of a temperate oligotrophic system similar to that of the Bermudas. The system alternates between two seasons: a warm and nutrient depleted summer during which the water-column is strongly stratified (from April to October), and a cold and nutrient rich winter during which the stratification of the water column breaks down (from November to March) (Fig. 2A and B). Second, we evaluate the changes occurring to the system when it is subject to a regime shift due to climate change, modeled as an increase of the average temperature and/or of the amplitude of the seasonal temperature variation over 100 years. The average temperature increases considered are +2, +4 and +10°C (applied uniformly over the water column), while the amplitude of the seasonal variation in surface temperatures is increased from 8°C to 18°C. These changes operate linearly over 100 years (Fig. 4A-C). Note that in order to artificially isolate the effect of regimes of temperature changes on the ecophysiology of the phytoplankton, the changes in temperature do not feed back onto the physics and chemistry of the water column: nutrient concentrations and the stratification profile of the water-column remain the same as temperature changes, while they would be expected to vary with temperature in an actual water-column. To assess whether such feedbacks could modify our predictions, we evaluated the effect of a drop in nutrient supply due to thermal stratification by decreasing the total nutrient mass in the system in some of the scenarios described above (see Discussion and Supplementary Material).

**Fig. 2:**
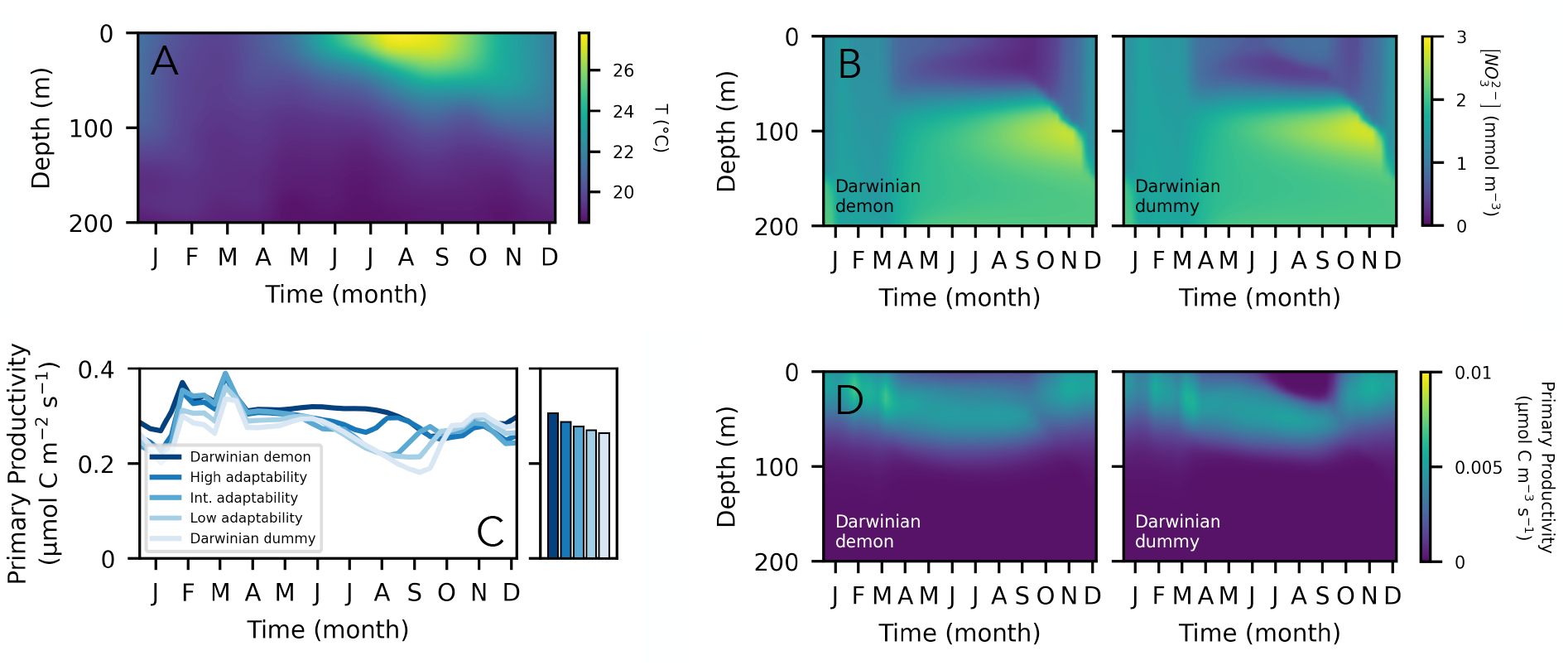
Simulated ecosystem dynamics in the Bermuda-like temperate oligotrophic system. The figure shows the seasonal variation of the vertical profile of the temperature (in °C) (A) and of the availability of nitrate (in μmol NO_3_^2−^ m^−3^) (B), and the seasonal variation of the primary productivity (PP) of the phytoplankton (in μmol C m^−3^ s^−1^) (C and D). The colored curves in panel C correspond to the vertical integration of the predicted PP in five evolutionary scenarios: Darwinian demon, high, intermediate, low mutation rate of the phytoplankton and Darwinian Dummy. The annual means are shown as colorbars on the right hand subpanel. In panels B and D, only the predictions obtained in the two extreme scenarios (i.e., the Darwinian Demon and Dummy scenarios) are shown.

**Fig. 3:**
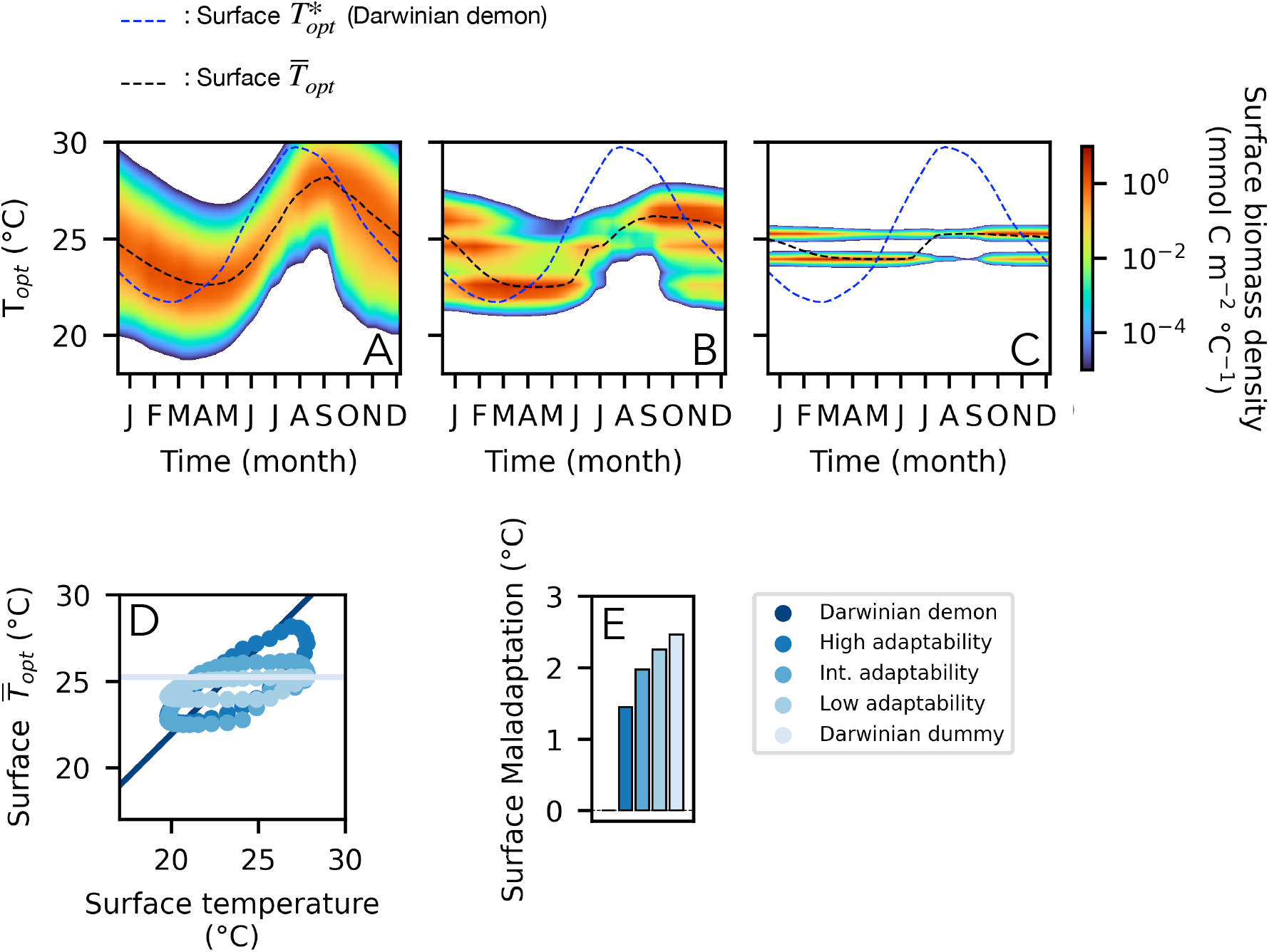
Adaptive response of the phytoplankton to the seasonal change in temperature at the surface. Panels A-C show the seasonal change in the trait distribution of the phytoplankton community in the first meters of the water-column when phytoplankton mutation rate is high, intermediate and low. The black dashed line corresponds to the community average thermal niche at the surface, 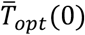, and the blue line to the evolutionary optimal trait 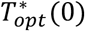 (i.e., corresponding to the trait of the perfectly adapting Darwinian demon; see Methods). Panel D shows, the match throughout the year between the average trait of the phytoplankton community at the surface 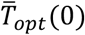 and the optimal trait 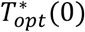 of the Darwinian demon given surface temperatures *T*(0) when the phytoplankton has a high, intermediate, and low mutation rate, and when the phytoplankton is a Darwinian dummy (no adaptation). For each of these evolutionary scenarios, the panel E shows the surface thermal maladaptation of the phytoplankton (measured as 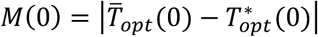) averaged over the year.

**Fig. 4:**
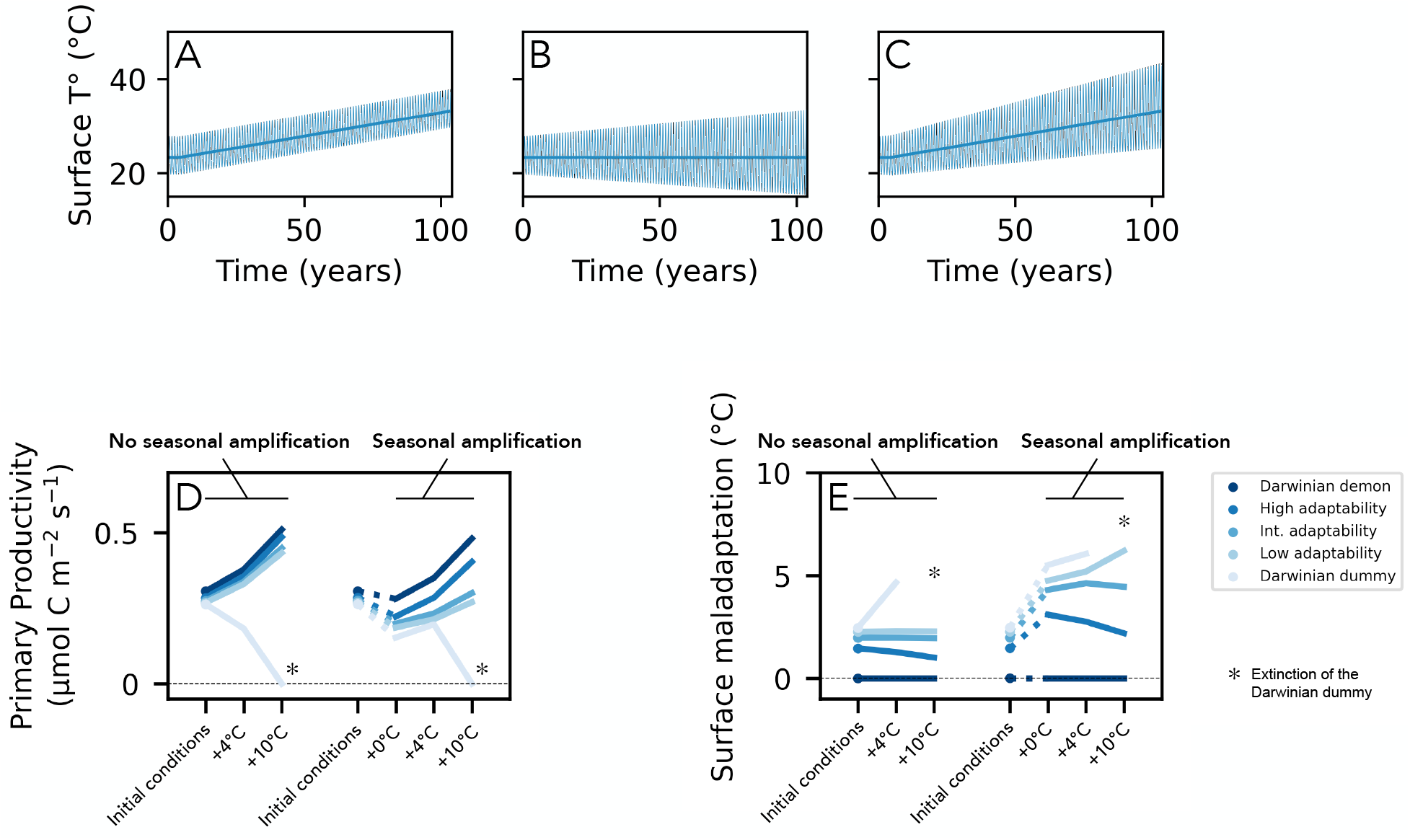
Simulated regimes of temperature change over 100 years (A-C) and phytoplankton primary productivity (D) and surface maladaptation (E) before and after these changes take place depending on the adaptive capacity of the phytoplankton. Climate change is here simulated either as an increase of the average temperatures of up to 10°C (A), as +10°C increase in the amplitude of the seasonal temperature variation (B) or as both simultaneously (C). Panels D and E show the primary productivity and surface maladaptation of the phytoplankton community in the initial conditions of the system and after a global temperature increase of +4 and +10°C coupled or not to an increase in seasonal amplitude. The * signs in panels D and E indicate when there has been an extinction of the phytoplankton population in the Darwinian dummy scenario.

### II.3. Evolutionary scenarios

We consider 5 evolutionary scenarios. The first one is the Darwinian demon scenario: there is only one type of phytoplankton, whose growth rate is always maximal (i.e., follows the Eppley envelope) for any given temperature. In other words, the thermal trait *T_opt_* of this Darwinian demon is always equal to the evolutionary optimum, noted 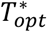, that maximizes growth given the local temperature (see supplementary materials and Fig. S1B on how we evaluate 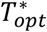). The second scenario is the Darwinian dummy: there is also only one population of phytoplankton, but its thermal niche is now fixed to make it well adapted to the systems’ average surface temperature. Then, in the three remaining scenarios, we run the eco-evolutionary SPEAD model considering a high, an intermediate, and a low value of mutation rate (*ν* = 10^−2^, 10^−5^, 10^−10^, respectively). We initialize the simulation with a phytoplankton community composed of a single phytoplankton population characterized by the same thermal niche as the Darwinian dummy in order for the mutational process to be the only source of the emergent functional diversity (the coexistence of multiple ecotypes with different thermal niches) in the phytoplankton community.

### II.4. Functional composition and maladaptation

At each depth *z* in the water column, we use the biomass-weighted average optimal growth temperature in the phytoplankton community, 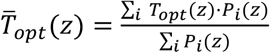, as an indicator of the dominant trait in the community. In the Darwinian demon scenario, 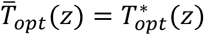, with 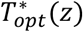 varying in the water-column following the environmental temperature, *T*(*z*) (see supplementary materials and Fig. S1B). In the Darwinian dummy scenario, 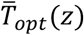 always corresponds to prescribed and fixed *T_opt_* of the phytoplankton community. The absolute difference between 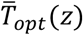 and the optimal value of the trait 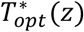 of the Darwinian demon (i.e., the value of *T_opt_* that maximizes growth given the temperature at *z*) is then used to evaluate the maladaptation *M*(*z*) of the phytoplankton community to its environmental conditions, with 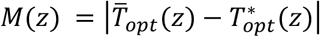. The unit of maladaptation is therefore the °C. Thereafter, we more specifically focus on the average trait and maladaptation of the phytoplankton at the surface (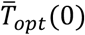 and *M*(0), respectively) where most of the primary productivity (PP) occurs.

## III. Results

### III.1. Eco-evolutionary response of the phytoplankton to seasonal changes

First, we simulate the seasonal dynamics at equilibrium of our temperate tropical ecosystem in the five scenarios described above: Darwinian demon (perfect and instantaneous adaptation), Darwinian dummy (no adaptation), and high, intermediate, and low mutation rates.

In the Darwinian demon scenario, biomass production is high during the late winter bloom and drops during the summer (Fig. 2C and D). The water column stratifies during the summer, the nutrient supply to the surface drops, and nutrients in the first 50 meters are depleted by phytoplankton consumption (Fig. 2B). Biomass production during the summer is then limited by the nutrient availability. In the Darwinian dummy scenario, the dynamics of biomass production is similar, except that the drop in productivity during the summer is much more pronounced (Fig. 2C and D). The biomass production in the Darwinian dummy scenario is approximately 15% lower than in the Darwininian demon scenario annually, and up to ~33% lower during the summer. In that case, the dip in production is not the result of a drop in nutrient availability as the nitrate concentration in the mixed layer increases as the summer progresses (Fig. 2B). Instead, it is due to surface temperature surpassing the fixed thermal tolerance of the Darwinian dummy, therefore restricted to deeper regions of the water column (~50 meters) where cooler temperatures prevail throughout the year (Fig. 2A and D). In other words, when the phytoplankton cannot adapt to temperature, the thermal niche of the phytoplankton is, in addition to nutrient availability, a determining factor of biomass production. This confinement of phytoplankton populations to their environmental (thermal) niche is called “environmental filtering” (Vallina et al. 2017). As higher adaptive capacities of the phytoplankton are considered, the phytoplankton is increasingly allowed near the surface, and PP is increasingly high during the summer relative to the Darwinian dummy (Fig. 2C): adaptation, allowing the phytoplankton thermal niche to shift with environmental temperatures, allows it to evade environmental filtering.

When the mutation rate of the phytoplankton is high (*ν* = 10^−2^), it adapts “on the fly” to seasonal changes in temperature (Fig. 3A). Near the surface, the distribution of the trait *T_opt_* in the community exhibits a single mode. The high mutation rate allows for a rapid succession of events of mutation and selection resulting in a relatively good match between community averaged *T_opt_* and the environmental temperature to which it is exposed (Fig. 3D and E). This fast-paced evolution allows a rapid recovery of PP after the initial dip as the summer progresses (Fig. 2C).

When the phytoplankton mutation rate is lower (*ν* = 10^−5^ and 10^−10^) the underlying mechanism to thermal adaptation is fundamentally different as it relies on successive shifts in relative abundance of ecotypes coexisting throughout the year, each being adapted to specific depths and periods of the year (Fig. 3B and C). In other words, the community averaged *T_opt_* mostly changes by competitive selection of the most adapted of a pool of pre-existing ecotypes rather than by the selection of new ecotypes appearing through trait mutation. In these scenarios, ecological selection thereby operates on a standing diversity that builds up over the years and is maintained by the spatio-temporal variability of the thermal conditions in the water-column. The lower the mutation rate is, the lower is the extent of the emerging standing diversity (i.e., of the variance of *T_opt_* in the community) on which selection can apply, and the higher is the mismatch between the average thermal niche of the phytoplankton and the temperature (Fig. 3D and E). When the phytoplankton mutation rate is the lowest (*ν* = 10^−10^), the range of the thermal adaptation becomes so narrow that there is almost no seasonal variability in the community averaged *T_opt_*. The phytoplankton community cannot therefore adapt to the warm temperature of the summer, nor to the cold temperature of the winter. The almost constant community averaged *T_opt_* corresponds to the annual average of the environmental temperature. As a result, PP throughout the year is very close to that of the Darwinian dummy scenario (Fig. 2C), yet this average “jack of all trades” *T_opt_* remains the best strategy possible given the limited adaptive capacity of the phytoplankton, allowing phytoplankton to survive the warm and cold extremes of the year.

To sum up, these results highlight that the capacity of the phytoplankton to adapt to seasonal changes in temperature can significantly influence its annual productivity (15% difference in annual PP between perfect and no adaptation), as thermal adaptation promotes the capacity of the phytoplankton to consume nutrients in the mixed layer of the water column throughout the year. Depending on the intensity of the mutational process, adaptation can rely either on a *de novo* diversity produced and selected upon as temperature changes, or on a standing diversity, emerging and maintained over the years. How will phytoplankton productivity change under those various eco-evolutionary regimes in a context of climate change?

### III.2. Eco-evolutionary response to global climate changes

Climate change is expected to manifest itself both through a secular trend of temperature increase and through more frequent extreme temperature events (Oliver et al. 2018, Frölicher et al. 2018, Oliver et al. 2019, Gruber et al. 2021, Burger et al. 2022). To simulate this two phenomena independently and combined, we change the temperature of the water-column by (*i*) imposing a temperature increase without an increase in seasonal amplitude, (*ii*) imposing an increase in seasonal amplitude without changing the average temperature, and (*iii*) imposing simultaneously an increase in average temperatures and in seasonal amplitude (see Methods; Fig. 4A-C). We look at the resulting changes in the annually averaged PP and in the annually averaged community averaged *T_opt_* and maladaptation of the phytoplankton community.

We first consider the consequences of temperature warming up over 100 years without changes in seasonal amplitude. In the Darwinian dummy scenario, PP after 100 years is systematically lower than its initial value, and the phytoplankton community actually collapses when the long-term temperature increase is the strongest (+10°C; Fig. 4D). As the average temperature increases, the phytoplankton community with a fixed (non-adaptive) thermal niche becomes increasingly unfit to near surface environmental conditions (Fig. 4E). During summer, environmental filtering results in phytoplankton being progressively confined to deeper regions of the water column, for longer periods of time. The phytoplankton eventually dies out when the resulting biomass loss becomes too large to be balanced out by biomass production during the winter bloom. By contrast, when phytoplankton individuals are Darwinian demons, a temperature increase results in an increase in PP. The extent of this boost in primary productivity is positively and linearly correlated to the amplitude of the temperature increase (+11%, +23% and +66% in PP for +2, +4 and +10°C respectively; Fig. 4D). According to the Eppley envelope, the metabolic activity of the phytoplankton accelerates with temperature. As temperature increases and because phytoplankton are able to adapt immediately and perfectly to it, its ability to produce biomass for a given nutrient influx increases (but see the Discussion section). When explicitly including adaptive evolution of phytoplankton in the model, the predicted changes in PP are very similar despite the large differences in the assumed mutation rates (Fig. 4D). As long as the mutational process is sufficient to sustain some degree of functional diversity, thermal adaptation can proceed. It would therefore appear that the Darwinian demon assumption is a robust approximation of the adaptive capacity of phytoplankton communities. In other words, thermal adaptation is very likely to be sufficiently rapid to keep up with global warming, even for a temperature increase as large as +10°C over 100 years. In fact, maladaptation remains constant over time at the surface in most scenarios and even drops when the mutation rate is high (Fig. 4E). This latter observation stems from the fact that phytoplankton individuals divide faster as temperature increases, generating more mutants, and the phytoplankton community is able to adapt faster to seasonal changes as a result. This is particularly true when thermal adaptation mostly relies on *de novo* diversity such as in the high mutation rate scenario (Fig. 3A).

We then consider the effect of seasonal amplification with no change in the annually averaged temperature (+0°C in the “seasonal amplification” case in Fig. 4). We observe that PP drops in every evolutionary scenarios, from −8% in the Darwinian demon scenario to −42% in the Darwinian dummy scenario (the other scenarios fall between these two extreme cases; Fig. 4D). The increased seasonality implies colder temperatures during the winter and warmer temperatures during the summer. As temperatures prevailing during the winter bloom get colder (Fig. 4A-C), the phytoplankton metabolism slows down (Fig. 1) and productivity lowers accordingly. This explains the productivity drop in the Darwinian demon scenario in spite of adaptation being perfect (Fig. 4D). Additionally, when adaptation is imperfect, the phytoplankton community progressively becomes unable to adapt to the increasingly large seasonal variation in temperature. This increased maladaptation drives productivity down (−22%, −28% and −31% for high, intermediate and low mutation rates respectively; Fig. 5B). Therefore, contrary to the previous case, the model predicts that mutation rate is a key parameter when increasing the seasonal amplitude without changing the average temperature (Fig. 4D).

**Fig. 5:**
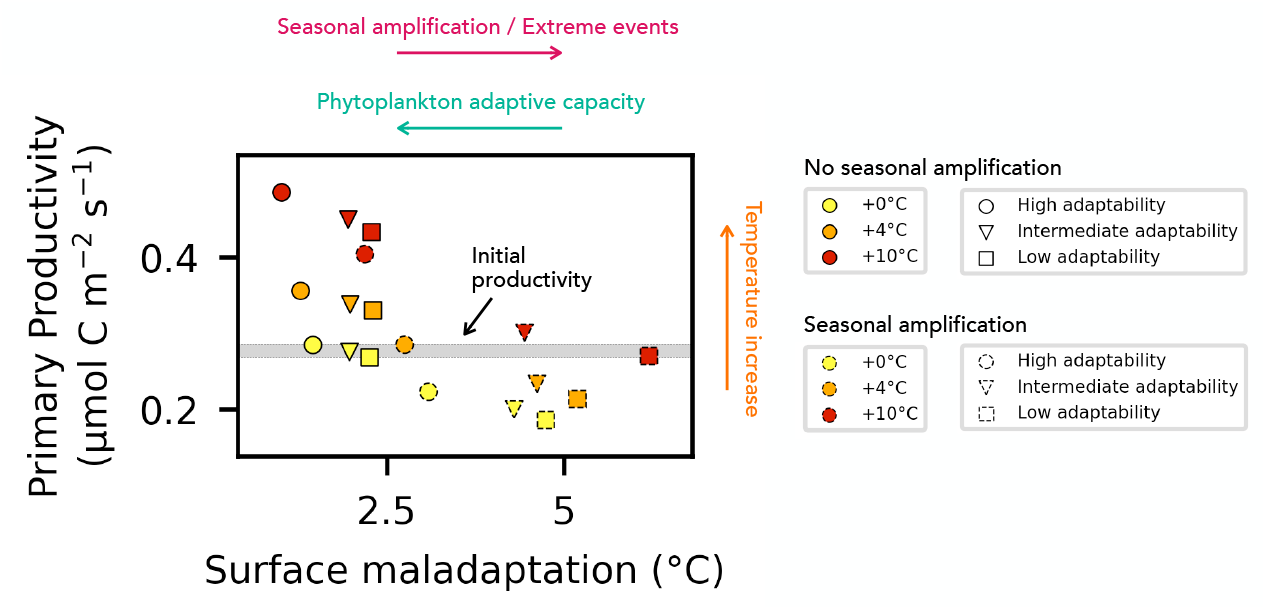
Link between primary productivity, temperature increase and maladaptation of the phytoplankton. Each dot corresponds to the state of the system after 100 years of a temperature increase of +0, +4 and +10°C coupled or not with seasonal amplification. We plotted only the scenarios in which the adaptive capacity of the phytoplankton is high, intermediate and low and excluded the Darwinian demon and dummy scenarios.

This is still the case when increasing temperatures and seasonality simultaneously. In the Darwinian demon scenario the negative effect of the seasonal amplification on PP is largely marginal compared to the positive effect of the increase in temperature (+3%, +14% and +57% for +2, +4 and +10°C respectively; Fig. 4D). However, this is not necessarily the case when adaptation is imperfect. A +10°C warming leads to PP increasing by 41% when the mutation rate is high, by 8% when it is intermediate, and by only 0.2% when the mutation rate is low (Fig. 4E); the phytoplankton still goes to extinction in the Darwinian dummy scenario. More importantly, for a +4°C warming, the prediction of increased PP is actually reversed (−0.8%, −15% and −20% for high, intermediate and low mutation rates; Fig. 4E)

To summarize, we find that primary productivity is expected to increase as global temperatures rise up in the context of a regime of climate change as long as phytoplankton communities are characterized by some capacity to adapt to temperature changes (Fig. 4D and 5). This effect could nevertheless be mitigated by an increase in thermal maladaptation (Fig. 4D-E and 5). Thermal maladaptation is expected to increase as short term variations of temperature become larger and faster (Fig. 5). The extent of this increased maladaptation then depends on the adaptive capacity of phytoplankton communities determined by their characteristic mutation rate (Fig. 4E). The predictions of our model are therefore especially sensitive to the adaptive capacity of the phytoplankton when increased seasonality is considered (Fig. 4D). As a result, our model predicts anything from a −31% to a +69% change in primary productivity depending on the assumed nature and amplitude of global warming, change in temperature seasonality, and extent of the phytoplankton adaptive capacity (Fig. 4D and 5).

## IV. Discussion

Using a 1D ocean model, we simulated the eco-evolutionary response of a phytoplankton community to long term and short term temperature changes, exploring a wide range of scenarios regarding the evolutionary properties of the phytoplankton –from perfect adaptation to no adaptation at all– and the regime of temperature changes. We found that when climate change is considered solely as a long-term trend of temperature increase, the adaptive response of phytoplankton is in most cases sufficiently rapid for phytoplankton to persist and even to see its biomass production increase, including for very large temperature increases (e.g, 10°C over 100 years; Fig. 4A). The commonly used Darwinian demon approximation then emerges as an acceptable representation of the adaptive capacity of the phytoplankton. However, when the amplitude of the seasonal variability of temperatures increases with the average temperatures, which is likely to occur in a regime of climate change (Oliver et al. 2018, Frölicher et al. 2018, Oliver et al. 2019, Gruber et al. 2021), the Darwinian demon approximation is no longer valid. We find that the efficacy of phytoplankton adaptation to increasingly extreme short-term temperature changes is indeed very dependent on the assumed phytoplankton mutation rate. The resulting maladaptation can substantially mitigate the prediction of increased biomass production. In some cases, when the amplitude of those short-term events is large compared to that of the long-term temperature increase, biomass production actually drops down as climate changes (Fig. 5). As primary production is one of the main driving forces of ocean biogeochemistry, our study highlights the importance of phytoplankton adaptation for the resilience of marine ecosystems to climate change. Our results are however obtained in an idealized framework, isolating the response of a local phytoplankton community to temperature changes alone. Several components of climate change and of the phytoplankton’s response to it are therefore neglected.

First, adaptation in the ocean is not a purely local process: when traits are locally selected, they can then be exported elsewhere through oceanic circulation and influence –positively or not– the phytoplankton adaptive response across the ocean (Leibold and Norberg 2004, Loeuille and Leibold 2008, Sauterey et al. 2017). Similarly, climate change will not just affect temperatures, but a multitude of other characteristics of oceanic systems (nutrient availability, pH, water stratification, horizontal circulation…) over “long” time scales (i.e., the century) but also through extreme compound events (Gruber et al 2021, Burger et al. 2022). We chose to evaluate in isolation the effect of temperature on the evolution of thermal niches and ignored those additional environmental stresses for simplicity. However, in natural systems under climate change, these environmental stresses could affect phytoplankton growth and adaptive capacity (Brennan et al. 2017). For instance, climate change is expected to enhance thermal stratification which should lead to a decreased nutrient supply to the euphotic zone, especially during summer. Climate change is also expected to increase depth of the mixed layer and the frequency and intensity of storms, which both tend to increase nutrient supplies by vertical turbulence (Sallée et al. 2021). Predicting how nutrient limitation will evolve under climate change is therefore challenging. Moreover, predicting the combined effect of nutrient limitations and temperature changes on phytoplankton growth and on the capacity of phytoplankton to adapt is a non-trivial issue. Empirical evidence suggests for instance that the effect of temperature changes on phytoplankton growth could be largely nullified in nutrient depleted conditions (Marañón et al. 2018). Although those questions are beyond the scope of this study, we provide some back-of-the-envelope estimates and additional discussion of how changes in nutrient supply affect our results (see Supplementary Material). Although the predicted increase in PP can be reversed by a concomitant decrease in nutrient supply, the main results regarding the sensitivity of PP evolution to the phytoplankton adaptive capacity are qualitatively unaffected. Those additional simulations actually show that the capacity of phytoplankton to adapt to temperature changes is diminished under more nutrient-limited conditions (see supplementary discussion). This suggests that such interacting effects between multiple environmental stressors might play a key role in determining the efficacy of phytoplankton adaptation across oceans (e.g., in nutrient depleted vs nutrient rich regions) and illustrates the need for further investigations. To explore these questions, we plan moving forward to implement our approach in a 3D model of ocean circulation. Such an integrated approach will allow evaluating in a spatially accurate environmental context how dispersal, the spatial distribution of environmental stressors and their covariance across oceans affect phytoplankton adaptation, and how this is expected to change in the context of climate change.

Additionally, several functional traits of phytoplankton communities are expected to evolve simultaneously in response to the combined selective pressures resulting from climate change (e.g., cell-size, traits related to nutrient, pH or light stresses). This raises an intriguing question: how does the simultaneous evolution of multiple traits influence the evolution of each singular trait (see Boyd et al. 2017 and Brennan et al. 2017 for some empirically grounded elements of response and Savage et al. 2007 for a first attempt at modeling multiple traits evolution)?

Similarly, phytoplankton are not the only marine organisms that climate change will affect. Higher trophic levels, characterized by a slower demography hence by a slower pace of evolutionary response will also be impacted by it. The same goes for the recycling microbial loop that determines how much of the organic matter circulating into marine ecosystems is recycled toward the ocean surface and how much is exported to the deep ocean (Cherabier and Ferriere 2022). Extending our approach to simulate the simultaneous evolution of multiple traits (as done in Le Gland et al. 2021) and the adaptive evolution of populations other than phytoplankton would provide an ideal modeling framework to address those currently unresolved issues.

Finally, although our results suggest that the adaptive capacity of phytoplankton might be a determining factor of the resilience of marine ecosystems to climate change, the problem of our very limited quantitative knowledge of what that capacity actually is still remains. Many macro-scale phenomena such as the emergence of the global biogeography of the thermal niches of the phytoplankton or the response of the phytoplankton to rapid climatic events such as El Niño/La Niña most likely involve, to some extent, thermal adaptation by the phytoplankton. Ocean-wide meta-genomic data sets are now becoming increasingly available (e.g., Sunagawa et al. 2020) and can be used to infer marine ecosystems’ composition and function (e.g., Chaffron et al. 2021) in such macro-scale contexts. We argue that constraining the parametrization of evolutionary approaches such as ours integrated in 3D models of ocean circulation based on the ability of these coupled models to reproduce those observed global scale phenomena will constitute an efficient way to quantify the adaptive properties of marine ecosystems, thereby providing a complementary approach to their experimental characterization.

## V. Concluding remarks

Our findings suggest that the resilience of marine ecosystems to climate change, and more specifically to the multiplication of extreme climatic events, will be determined by the ability of phytoplankton communities to adapt. Current Earth System Models, most of which assume perfect and instantaneous thermal adaptation by the phytoplankton, are therefore likely to overestimate this resilience. While the present exploratory work shows how our limited quantitative knowledge of phytoplankton adaptive properties limits our ability to predict the resilience of marine ecosystems to climate change, it also paves the way toward improving that knowledge through the use of models of phytoplankton evolution. Ultimately, we think that approaches such as ours may play a key role in increasing the accuracy of Earth System Models’ climate projections.

## Acknowledgement

This research is part of the GOMMA project founded by the CSIC – IEO (grant number PID2020-119803GB-I00).

