## Supplementary materials for "Phytoplankton adaptive resilience to climate change collapses in case of extreme events – A modeling study"

### I. Thermal niche of the phytoplankton

#### I.1. Parametrization

Phytoplankton growth $g$ is described in the model as follows

$$\text{(E1)} \quad g = g_{max} \cdot \gamma_T \cdot \gamma_N \cdot \gamma_I \cdot \gamma_{p_{CO_2}} \cdot \gamma_P$$

where $g_{max}$ is the maximum growth rate of phytoplankton individuals and $\gamma_T$, $\gamma_N$, $\gamma_I$, $\gamma_{p_{CO_2}}$ and $\gamma_P$ the growth limitation factors associated with temperature, with the availability of nutrients, light, and dissolved $CO_2$, and with the Allee effect (see discussion below), respectively. The term $\gamma_T$ corresponds to the thermal reaction norm of phytoplankton individuals (Fig. 1 and S1A). Following Le Gland et al. 2021, the thermal reaction norm $\gamma_{T,i}$ of a phytoplankton population $P_i$, with a temperature of optimal growth $T_{opt,i}$ is expressed as:

$$\text{(E2)} \quad \begin{aligned} &\gamma_{T,i} = e^{\alpha(T_{opt,i}-T_{ref})} \cdot e^{\frac{T-T_{opt,i}}{\omega}} \cdot \frac{(T_{opt,i}+\omega-T)}{\omega} \text{ when } T - T_{opt,i} < \omega, \\ &\gamma_{T,i} = 0 \text{ when } T - T_{opt,i} > \omega. \end{aligned}$$

Each term corresponds to one specific aspect of the temperature dependence of phytoplankton growth (Fig. S1A). The first term describes how the maximum growth of the population $i$, achieved when $T = T_{opt,i}$, increases with the value of $T_{opt,i}$, following the Norberg-Eppley model, with $\alpha$ and $T_{ref}$ being the exponent and reference temperature of the Eppley envelope (Le Gland et at. 2021). The second term describes how growth progressively converges toward 0 when temperatures get lower than $T_{opt}$. The third term describes how growth abruptly drops to zero when temperatures get higher than $T_{opt}$. In those two terms, $\omega$ corresponds to the thermal tolerance of the phytoplankton population. It determines how fast phytoplankton growth converges toward zero at low temperatures, and what is the higher temperature limit for which growth is superior to zero. In our simulations, we used a thermal tolerance of 5°C (which corresponds to the thermal reaction norms plotted in Fig. 1 and S1).

#### I.2. Evolutionary optimal thermal trait

For any given temperature $T$ of the system, there is a trait $T_{opt}^*$ that maximizes phytoplankton growth (Fig. S1B). $T_{opt}^*$ is therefore an evolutionary optimum. By definition, the trait of the Darwinian demon is always equal to $T_{opt}^*$. Also by definition, when $T_{opt} = T_{opt}^*$ then $\frac{d\gamma_T(T)}{dT_{opt}} = 0$. From (E2) we get the expression of $\frac{d\gamma_T(T)}{dT_{opt}}$:

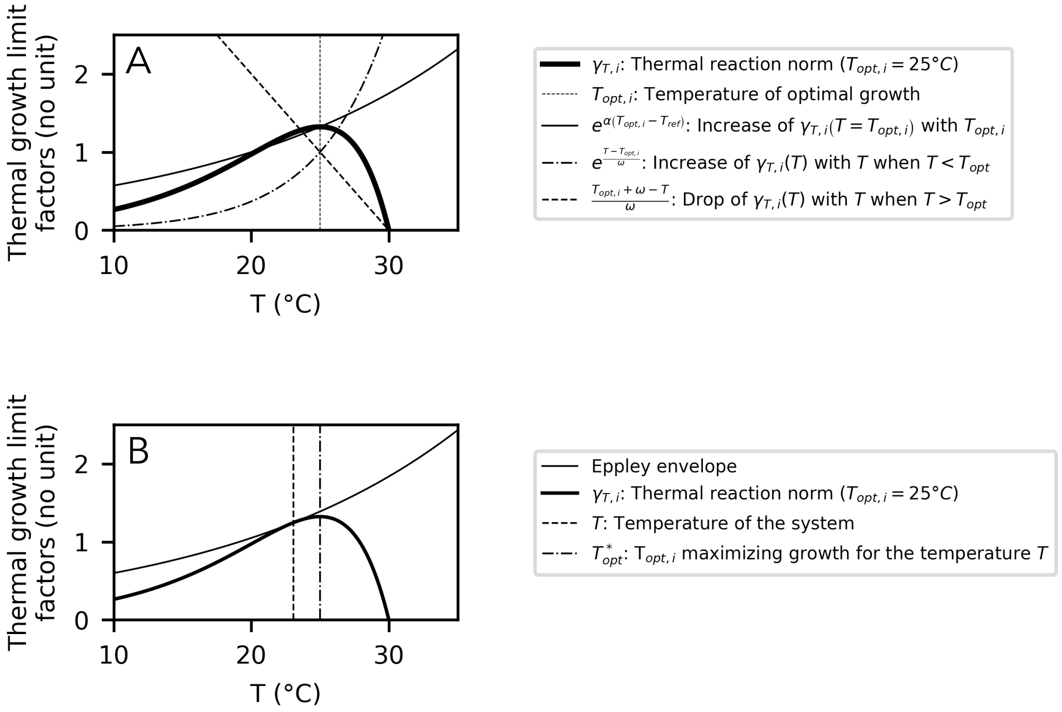


**Supplementary Figure 1: Temperature dependence of phytoplankton growth.** Panel A shows how the combination of three temperature dependent terms determines the shape of the thermal reaction norm of a phytoplankton population $P_i$ characterized by a $T_{opt,i}$ = 25°C. Panel B shows how, for a given temperature $T$, growth is maximized by a phytoplankton population characterized by a $T_{opt} = T^*_{opt} \approx T + 1.94$°C.

$$\text{(E3)} \quad \frac{d\gamma_T(T)}{dT_{opt}} = \frac{\left(T - T_{opt} + \alpha\omega\left(T_{opt} - T + \omega\right)\right)e^{\frac{T - T_{opt} + \alpha\omega\left(T_{opt} - T_{ref}\right)}{\omega}}}{\omega^2}.$$

Solving $\frac{d\gamma_T(T)}{dT_{opt}} = 0$ is therefore equivalent to solving $\left(T - T_{opt} + \alpha\omega\left(T_{opt} - T + \omega\right)\right) = 0$. This equation accepts the solution

$$\text{(E4)} \quad T^*_{opt} = T + \frac{\alpha\omega^2}{1 - \alpha\omega}.$$

With the parametrization used in our simulation, $T^*_{opt} \approx T + 1.94$ °C.

**Supplementary Table 1: temperature dependence of phytoplankton growth**

| **Parameter** | **symbol** | **value** | **unit** |
|---|---|---|---|
| Eppley exponent | $\alpha$ | 0.056 | $°C^{-1}$ |
| Eppley reference temperature | $T_{ref}$ | 20 | °C |
| Thermal tolerance | $\omega$ | 5 | °C |

### II. Eco-evolutionary model

#### II.1. Discrete trait-diffusion method

The model solves the population dynamics of 50 phytoplankton "subtypes". Each of these subtypes $P_i$ is characterized by a specific thermal niche and by a specific temperature of optimal growth $T_{opt,i}$, ranging from 18°C to 50.34°C by increment of 0.66°C. We refer to the ensemble of these populations as the phytoplankton community, and to the relative abundance of those subtypes as the functional composition of the phytoplankton community. The net growth rate of these phytoplankton populations is expressed as:

$$\text{(E5)} \quad s_i = (g_i - d)P_i$$

where $g_i$, the individual growth rate of the population $P_i$, depends on the adequation of the trait of the population $T_{opt,i}$ and the temperature of the system $T$ according to equations (E1) and (E2) and where $d$ is mortality. This net growth rate corresponds to the fitness of each of the 50 populations. Additionally, the model describes how, at each generation, a proportion of the offspring produced by each population is carrying phenotypically altering mutations that results in their trait being different from that of their ancestors. The effect of this mutational process on the functional composition of the phytoplankton community is simulated as a diffusion process (Van Der Laan and Hogeweg 1995, Leimar et al. 2008, Sauterey et al. 2017, Le Gland et al. 2021) as follows:

$$\text{(E6)} \quad \frac{dP_i}{dt} = s_i + \nu \cdot \sigma^2 \frac{\partial^2 g_i P_i}{\partial {T_{opt}}^2}$$

where $\nu$ is the probability of a mutation occurring (i.e., the mutation rate) at each division event, $\sigma$ the average phenotypic effect of this mutation (here considered to be equal to one), and $\frac{\partial^2 g_i P_i}{\partial {T_{opt}}^2}$ the local gradient (around $T_{opt,i}$) of the individual division rate along the dimension of the trait $T_{opt}$. The second term of this expression therefore describes the mutational flux of individuals in and out of the population $P_i$. By considering 50 phytoplankton subtypes – hence a finite number of 50 trait values – instead of considering the fate of individuals carrying every possible values of $T_{opt}$, we actually perform a discrete approximation of the trait space. In the context of this approximation, the equation (E6) becomes

$$\frac{dP_i}{dt} = s_i + \nu \cdot \sigma^2 \frac{g_{i+1}P_{i+1} + g_{i-1}P_{i-1} - 2g_iP_i}{\Delta {T_{opt}}^2},$$

$$\text{(E7)} \quad \frac{dP_i}{dt} = s_i + \nu \cdot \sigma^2 \frac{g_{i+1}P_{i+1} - g_iP_i}{\Delta {T_{opt}}^2} \text{ when i = 0 and}$$

$$\frac{dP_i}{dt} = s_i + \nu \cdot \sigma^2 \frac{g_{i-1}P_{i-1} - g_iP_i}{\Delta {T_{opt}}^2} \text{ when i = 50.}$$

with $\Delta T_{opt} = 0.66$°C.

#### II.2. Allee effect

One of the known drawbacks of that method of simulating eco-evolutionary process (Perthame and Gauduchon 2009, Sauterey et al. 2017) is that the diffusion terms tend to result in the emergence of trait distributions characterized by infinitely long tails of infinitesimal populations.

Although it is not a problem when considering a biological system under continuous selective pressure, it can become one when considering a fluctuating ecosystem such as a seasonal oceanic ecosystem. Those infinitesimally rare populations can then take over the ecosystem after a brusque change in the environmental conditions favorable to them, thus generating unrealistic evolutionary jumps. To solve this problem, we implement into our model of phytoplankton growth an Allee effect $\gamma_P$ (from the seminal work of Allee et al. 1949) which describes how the fitness of the individuals of the population $P_i$ drops with the population abundance:

$$\text{(E8)} \qquad \gamma_P = \frac{P_i}{P_i + A}$$

where $A$ is the Allee constant such that when $P_i = A$ the individuals' growth rate is half of their maximal growth rate. This limiting factor results in the population growth rapidly dropping toward 0 when $P_i << A$. Consequently, a rare population can only take over the ecosystem after an environmental change only if it has been maintained in the system at sufficient levels of abundance, either by mutation, or by vertical mixing.

### III. Interacting effects of nutrient availability and temperature change on thermal adaptation

In the context of climate change, phytoplankton populations are expected to face multiple environmental stresses in addition to long-term and short-term temperature changes. It remains unclear how the combination of these multiple environmental stresses will affect the physiological and adaptive responses of phytoplankton individuals and communities to every individual stress (Boyd et al. 2017, Brennan et al. 2017, Marañon et al. 2018). As mentioned in the core of this study, we artificially isolated the effect of temperature on phytoplankton from any other environmental stresses. Would our predictions change should other environmental

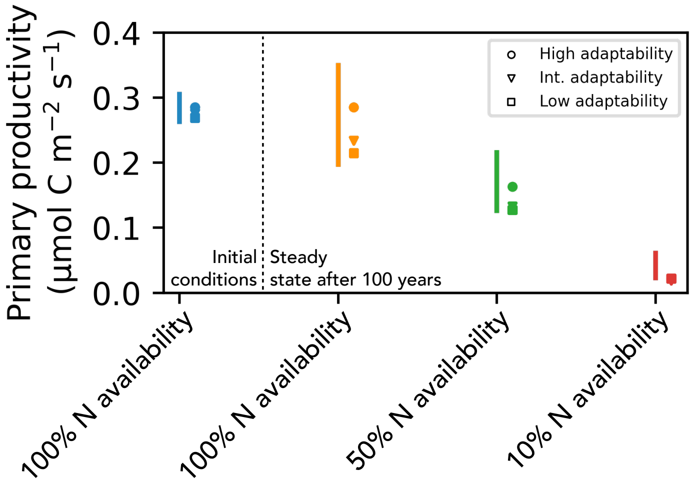


**Supplementary Figure 2: Ranges of predicted surface primary productivity as a function of nitrogen (N) availability in the water column.** The blue, orange, green and red vertical lines correspond to the ranges of predicted primary productivity prior to climate change and after 100 years of climate change without reduction of the nutrient availability, with nutrient availability being reduced by a factor 2, or by a factor 10, respectively. The climate change consists of +4°C increase in average temperature and a +10°C increase in temperature annual variability. The minimum value of the range corresponds to the Darwinian dummy scenario, the maximum value to the Darwinian demon scenario. The circles, triangles and squares correspond to the scenarios of high, intermediate, and low adaptive capacity of the phytoplankton respectively.

stresses be accounted for? In a first attempts at answering this question, we ran new sets of simulations in which, in addition to an average temperature increase of 4°C and an increase in seasonal variation of temperature of 10°C, the phytoplankton is also exposed to a reduction of the nutrient availability by a factor 2 and 10. Contrary to the regime of temperature change, this reduction of the nutrient availability takes place at the beginning of the simulation, not progressively throughout the 100 years of the simulation. The results are shown in Figure S2.

First, we find that a decrease in nutrient availability can counteract the effect of temperature increase on phytoplankton growth and can break down the prediction of increased primary productivity in a regime of climate change. Second, we find that although the relative spread of the prediction range remains of the same order regardless of nutrient availability (i.e., the minimum predicted value of primary productivity is lower than the maximum by 43, 41 and 62% when nutrient availability is unchanged, divided by 2 and divided by 10, respectively), the absolute spread of the predictions drops as nutrient availability is decreased. Finally, we find that the adaptive capacity of the phytoplankton increasingly diverges from the Darwinian demon scenario as the nutrient availabilities considered are smaller: nutrient depleted conditions result to lower division rates which in turn results in a decreased rate of adaptation by the phytoplankton. We see this last result as particularly interesting as it highlights that adaptation of specific functional traits (here the thermal niche) might be influenced by apparently unrelated environmental stressors (here nutrient availability) and suggests that studying such interacting effects in the more realistic context of a 3D circulation model might be key to better predict the overall adaptive response of phytoplankton communities to multi-faceted environmental changes.
